# Phenotypic Covariation And Morphological Diversification In The Ruminant Skull

**DOI:** 10.1101/017533

**Authors:** Annat Haber

## Abstract

Differences among clades in their diversification patterns result from a combination of extrinsic and intrinsic factors. In this study I examined the role of intrinsic factors in the morphological diversification of ruminants in general, and in the differences between bovids and cervids in particular. Using skull morphology, which embodies many of the adaptations that distinguish bovids and cervids, I examined 132 of the 200 extant ruminant species. As a proxy for intrinsic constraints I quantified different aspects of the phenotypic covariation structure within species, and compared them with the among-species divergence patterns, using phylogenetic comparative methods. My results show that for most species, divergence is well aligned with their phenotypic covariance matrix, and those that are better aligned have diverged further away from their ancestor. Bovids have dispersed into a wider range of directions in morphospace than cervids, and their overall disparity is higher. This difference is best explained by the lower eccentricity of bovids’ within-species covariance matrices. These results are consistent with the role of intrinsic constraints in determining amount, range, and direction of dispersion, and demonstrate that intrinsic constraints can influence macroevolutionary patterns even as the covariance structure evolves.

## Introduction

Diversity is distributed unevenly across the tree of life. This unevenness is the result of both extrinsic factors, such as environmental changes and biotic interactions, and intrinsic factors, such as developmental and genetic interactions. While extrinsic factors provide the opportunities for diversification, intrinsic factors determine the variation available for natural selection to work on, and therefore the potential of the population to take advantage of these opportunities. Studying this potential is therefore crucial for understanding macroevolutionary patterns. A common approach for studying the effect of intrinsic factors is to look at the structure of phenotypic and genetic (co)variation within the population, and compare it to patterns of divergence among populations (e.g., Lande 1979; Zeng 1988; Armbruster 1996; Schluter 1996; Arnold et al. 2001; Ackermann and Cheverud 2002; Hansen and Houle 2008; Hohenlohe and Arnold 2008; Klingenberg 2008; Marroig et al. 2009; Bolstad et al. 2014).

According to the quantitative genetics framework (Lande 1979), the rate and direction of short-term evolution depends on the relative alignment and magnitude of selection and the within-population genetic (**G**) and phenotypic (**P**) covariation. A closer alignment between the major axis of covariation and selection allows the population to evolve more rapidly in the direction of selection because more variation is available in that direction. A greater angle between them will divert the population further away from the optimum, slowing down its evolution. A direct extension of this framework to the macroevolutionary scale predicts that the divergence of populations will initially be biased by the major axis of their ancestral covariation structure, resulting in a close alignment between the within-population covariation and the among-population divergence (Zeng 1988; Armbruster 1996; Schluter 1996). This bias is expected to diminish with time. Hansen and Houle (2008) expanded this approach to quantify different aspects of the evolutionary potential and provide a null expectation for evaluating the extent of the bias (also Hansen and Voje 2011; Bolstad et al. 2014). They defined the evolvability of a single trait as the expected change in trait value in response to directional selection of unit strength per generation, measured as the mean-scaled trait variance. In a multivariate space, evolvability is defined with respect to a specific direction; e.g., that in which selection is acting or that in which divergence has occurred (*e*(**d**_SP_) and *e*(**d**_CL_) in table 1). The conditional evolvability is defined as the expected change in a given direction assuming stabilizing selection in the remaining directions, thus accounting for trait covariances as well as variances. When averaged across random directions these measures provide the null expectation of no bias (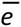 and 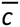 in table 1).

**Table 1.** Parameters and metrics included in this study, their symbols, and their interpretations

| Symbol | Measure |  | Meaning |
| --- | --- | --- | --- |
| **P** | Pooled within-population phenotypic V/CV matrix; based on individuals and corrected for sex and subspecies mean |  | Variation and covariation of traits among individuals within a species |
| **P<sub>AV</sub>** | Average pooled within-population phenotypic V/CV matrix; based on individuals and corrected for species mean, as well as sex and subspecies |  | Variation and covariation of traits among individuals within a species, averaged across species within a clade. |
| **D<sub>IC</sub>** | Among-population phenotypic V/CV matrix; based on species means and corrected for phylogenetic correlation assuming BM (i.e., V/CV of independent contrasts; Revell 2007) |  | Evolutionary rate matrix; rate of evolution and coevolution of traits among species within a clade |
| $\lambda$ | An eigenvalue of **P** or **D<sub>IC</sub>** | | |
| $\beta$ | A vector of $p$ elements, drawn randomly from a normal distribution with mean of zero and SD of one, normed to a unit length | | Selection gradient; direction of selection operating on each trait independently of other traits |
| Average evolvability $\bar{e}$ | The average mean-scaled phenotypic variance in **P**;<br>Proportional to matrix size | $E\left[\beta^t \mathbf{P} \beta\right]$ or $E\left[\lambda(\mathbf{P})\right]$ | The expected ability of the population to respond to selection in a random direction independently of other directions |
| Average conditional evolvability $\bar{c}$ | The average mean-scaled phenotypic variance in **P**, accounting for trait covariances;<br>Includes both matrix size and matrix shape | $E\left[\left(\beta^t \mathbf{P}^{-1} \beta\right)^{-1}\right]$ | The expected ability of the population to respond to selection in a random direction, assuming stabilizing selection in other directions |
| Average flexibility $\bar{f}$ | Cosine of the angle between $\beta$ and the response vector, averaged over random vectors;<br>Matrix shape only | $E\left[\frac{\beta^t \mathbf{P} \beta}{\sqrt{\Sigma(\mathbf{P} \beta)^2}}\right]$ | The expected ability of the population to track in the direction of selection; reflecting the range of random directions into which the population can potentially evolve with relative precision |
| Within-pop. Eccentricity rSDE(**P**) | Relative standard deviation of the eigenvalues of **P**<sup>a,b</sup>;<br>Matrix shape only | $\sqrt{\frac{p \sum\left(\lambda_i-\bar{\lambda}\right)^2}{(p-1) \sum \lambda \sum \lambda}}$ | The relative ability to track in different directions; reflecting the range of orthogonal directions into which the population can potentially evolve with relative precision |
| Among-pop. eccentricity rSDE(**D<sub>IC</sub>**) | Relative standard deviation of the eigenvalues of **D<sub>IC</sub>** |  | Eccentricity of dispersion; the range of directions in space into which the species have diverged |
| **d<sub>CL</sub>** | An eigenvector of **D<sub>IC</sub>** |  | Direction of clade’s dispersion |
| **d<sub>SP</sub>** | Difference vector between a species mean and its clade’s phylogenetically-weighted mean (inferred root state) assuming Brownian motion, standardized by that mean and normed to a unit length. |  | Direction of morphological divergence between a species and its clade’s ancestral state |
| $d^2$ | Squared distance between a species mean and its clade’s inferred root | $\mathbf{d}_{\text{sp}}^t \mathbf{d}_{\text{sp}}$ | Amount of morphological divergence between a species and its clade’s ancestral state |
| Disparity | Average distance between species means and the clade’s inferred root<sup>c</sup> | $\frac{\sum d^2}{\left(N_{\text{sp}}-1\right)}$ | Total amount of clade’s dispersion |
|  | Trace of **D<sub>IC</sub>** |  | Rate of dispersion |
| $e\left(\mathbf{d}_{\text{sp}}\right)$ | Mean-scaled phenotypic variance in the direction of species divergence | $\mathbf{d}_{\text{sp}}^t \mathbf{P}_{\text{av}} \mathbf{d}_{\text{sp}}$ | The variation available for species, on average, in the direction of their divergence |
| $c\left(\mathbf{d}_{\text{sp}}\right)$ | Mean-scaled phenotypic variance in the direction of species divergence, accounting for trait covariances | $E\left[\frac{1}{\mathbf{d}_{\text{sp}}^t \mathbf{P}_{\text{av}}^{-1} \mathbf{d}_{\text{sp}}}\right]$ | The variation available for species, on average, in the direction of their divergence assuming stabilizing selection in other directions |
| $e\left(\mathbf{d}_{\text{cl}}\right)$ | Mean-scaled phenotypic variance available along the eigenvectors of **D<sub>IC</sub>** | $\mathbf{d}_{\text{CL}}^t \mathbf{P} \mathbf{d}_{\text{CL}}$ | The variation available for each species in the direction of their clade’s diversification |

<sup>a</sup> *p* is number of traits<sup>b</sup> Developed by Van Valen (1974) and is suitable for both covariances and correlations. For correlations, it reduces to rSDE given in Pavlicev et al. (2009). For covariances, it standardizes for total variance thus factoring out matrix size.<sup>c</sup> *N<sub>SP</sub>* is number of species

For both the micro- and the macro-evolutionary scales, this framework focuses on the relative alignment between covariation and selection; it measures the amount of variation available in a specific direction and therefore the potential to evolve faster in that direction. However, as the phylogenetic scale and time span of the study increase it becomes more difficult to reconstruct relative alignment of selection and covariation at any given time, and more likely that either or both have changed repeatedly, so that their time-averaged estimation (or ancestral reconstruction) becomes less informative. The stability and predictability of **P** and **G** has been a particularly thorny issue when attempting to bridge the micro and macro scales. It is now widely agreed that the relevant question is when and how they evolve rather than if, and that the answer is complex and context-dependent (Arnold et al. 2008; Haber 2015; and references therein). Moreover, in the long run, selection is more likely to shift often enough and in directions different enough to present the population with a diverse array of challenges and opportunities (Vermeij 1973, Jones et al. 2012). Therefore, we might expect macroevolutionary patterns to be largely determined – and better predicted – by species’ ability to respond effectively to a wide range of selective pressures (Vermeij 1973; Liem 1973; Draghi and Wagner 2008) rather than their ability to respond faster in any particular direction. Regardless of scale, understanding the potential of populations to respond to whatever may come is becoming increasingly relevant nowadays with global environmental changes becoming more drastic and less predictable (Kopp and Matuszewski 2014).

I consider here four measures that could reflect the potential to respond to a wide range of selective pressures: 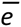, 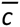, 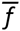, and rSDE(P) (see table 1 for details). The average evolvabiliy (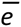) and conditional evolvabiliy (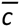) developed by Hansen and Houle (2008) measure the variance available in any random direction on average, and thus the average potential to respond to any selection vector. A higher value of 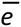 or 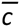 can be interpreted as a greater potential to respond to a wider range of selection vectors, where 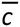 assumes stabilizing selection in the remaining directions. Both are proportional to total variance (i.e., matrix size), and 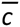 accounts also for trait covariances (i.e., matrix shape). Another measure based on Hansen and Houle (2008) is the average flexibility (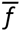; Marroig et al. 2009; Rolian 2009), which measures how well the population response is aligned with the direction of selection, on average. A higher value means that the population is less biased by its major axis of covariation and is able to align more closely with a wider range of selection vectors (i.e., more flexible). 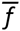 quantifies matrix shape only, not accounting for the orientation and size of the matrix.

Another metric that quantifies only matrix shape is the relative standard deviation of the eigenvalues (rSDE(**P**) in table 1). This metric, in its various forms, has been referred to as integration level in studies of morphological integration (Wagner 1984, Cheverud et al. 1989, Pavlicev et al. 2009, Haber 2011). Its use as a measure of integration has been criticized recently for being merely descriptive with no link to the theory of integration and evolution (Armbruster et al. 2014). However, here I use it to quantify matrix shape – as originally suggested by Van Valen (1974) – and reinterpret it in the context of Hansen and Houle’s (2008) model. Since evolvability is defined as the (mean-scaled) variance in a given direction, the eigenvalues of a matrix measure the evolvabilities along its eigenvectors. Therefore, rSDE(**P**) measures the variation in the ability to respond in different (orthogonal) directions. Because rSDE(**P**) is standardized by the total variance of the matrix, it captures matrix shape only. When the variation in the sample is distributed more evenly in different directions, matrix shape is less eccentric, and rSDE(**P**) is lower. By definition, matrices with a lower eccentricity are less biased by their major axis of covariation. Therefore, we would expect rSDE(**P**) to have a tight negative correlation with 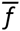, as indeed was found by Marroig et al. (2009) and Goswami et al. (2014) for mammals and by Hopkins et al. (unpublished data) for fiddler crabs. Simulations indicate that this relationship persists for a wide range of matrices that differ in their number of variables, mean correlation, and heterogeneity (see online appendix B; also Goswami et al. 2014). To the extent that the observed covariation matrix reflects underlying integrating factors, a more eccentric matrix implies a more strongly integrated body-plan. However, the link between rSDE(**P**) and integration is not the focus of this study. Instead, I focus here on the association between rSDE(**P**) and macroevolution.

Because 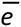 and 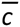 are proportional to matrix size (i.e., total variance), a direct extrapolation from micro- to macroevolution assumes a linear relationship between the probability of responding effectively and the variance available to selection at any given time. In this context, ‘responding effectively’ means reaching close enough to the optimum to avoid extinction or competitive displacement. We might, however, imagine a situation where the relationship is not linear, involving a threshold of variance that needs to be exceeded in order to avoid extinction and diverge successfully (Gomulkiewicz and Houle 2009, Jones et al. 2012, Kopp and Matuszewski 2014). When the variation is distributed more evenly across the different dimensions, there would be more directions in which the threshold is likely to be exceeded, for a given matrix size (Klingenberg 2005; Jones et al. 2012). Therefore, a population with lower eccentricity would be able to respond effectively in more directions than a population with higher eccentricity and the same matrix size. By extension, because of the threshold, clades whose species maintain lower eccentricity might show higher disparity in morphospace as well as a more even dispersion.

Which variational property best predicts macroevolutionary patterns is still an open question (Hansen 2012; Bolstad et al. 2014). Each approach quantifies different aspects of the evolutionary potential and carries a different set of assumptions, particularly regarding the heterogeneity and predictability of past and future selection, as I discuss further below. Under what conditions the different assumptions are warranted and the different aspects become more relevant, is still largely unknown. Here I present an empirical example using the ruminant skull that provides some insights to these questions.

Ruminants are especially suitable for understanding how variational properties of populations contribute to differences in diversity among higher clades. Ruminants comprise the bulk of Artiodactyla (Vrba and Schaller 2000; Theodor et al. 2005), and consist of Tragulidae (chevrotains) and Pecora (giraffes, pronghorn, musk deer, bovids, and cervids). Bovids and cervids are currently the most species-rich families within Ruminantia (143 and 51 extant species, respectively). Yet, their taxonomic richness is not too high to be reasonably sampled within the scope of this study, allowing for both good taxonomic coverage and large within-species samples. In addition, their phylogenetic history is relatively well understood (Hernández Fernandez and Vrba 2005; Price et al. 2005; Gilbert et al. 2006; Marcot 2007; Agnarsson and May-Collado 2008; Decker et al. 2009), and their fossil record is exceptionally good for vertebrates (Geist 1985, 1987; Janis 1989; Vrba and Schaller 2000; Marcot 2004; Pitra et al. 2004; Ropiquet and Hassanin 2005; Theodore et al. 2005; Gillbert et al. 2006; Janis 2008; Heywood 2010). Previous studies have provided ample evidence that ruminants have faced a wide range of selective pressures due mostly to global changes such as Plio-Pleistocene glacial cycles and the opening of grasslands. At the same time, the fossil record reveals that cervids have had about the same opportunities as bovids to exploit open habitats and arid grasslands at least since the late Miocene (Janis 2008; Heywood 2010). Yet, no cervid has evolved into a true grazer. Cervids are mostly browsers, occupying habitats with dense vegetation, whereas bovids exhibit a high diversity of feeding adaptations and a wider range of body size and shape (Allard et al. 1992; Spencer 1997; Sinclair 2000; Marcot 2004; Janis 2008; Heywood 2010). These differences have left researchers puzzled as to why cervids have not taken advantage of the same opportunities as bovids (Janis 2008; Heywood 2010). In this study I ask whether properties of the within-population covariation structure could have contributed to the morphological diversification of ruminants in general, and to the contrast between bovids and cervids in particular.

## Materials and Methods

### Data collection and preparation

A total of 2857 skulls were included in this study, representing 5 out of 8 species of tragulids, 3 out of 7 extant species of moschids, 34 out of 51 extant species of cervids, and 87 out of 143 extant species of bovids. All 19 extant cervid genera are represented except *Przewalskium albirostris,* which has been endangered for the last few decades. Of the 50 extant bovid genera, 40 are represented. The remaining 10 consist of only 1 to 3 species each and are all nested within otherwise well-represented clades. All domesticated species were excluded. Taxonomic assignments of specimens were standardized following Grubb (2002). Only prime age adults were measured, as determined by their dentition and cranial sutures. Three-dimensional coordinates were recorded for 43 landmarks using MicroScribe MLX. Landmark definition was based on the standard measurements recommended by von den Driesch (1976), as well as other studies of the artiodactyl skull (Janis 1990; Mendoza et al. 2002; Semprebon et al. 2004). All landmarks except those adjacent to teeth are based on suture junctions (online table A1). Data processing involved unifying the dorsal and ventral aspects of the skull, averaging the left and right, and identifying outliers (see online Appendix A for details). Variances due to measurement error were found to be between 1 to 3 orders of magnitude lower than the inter-specimen variance (see online Appendix A).

Two datasets of interlandmark distances were created. For the first set (‘ILtes’), 107 interlandmark distances were defined based on Delaunay tessellation of the symmetric mean configuration of all specimens, function *delaunayn* in the R package *geometry* (Barber et al. 2014). This procedure maximizes coverage while minimizing redundancy and crossing over spatial modules. The second set (‘IL32’) included 32 interlandmark distances selected based on comparability with other studies (e.g., von den Driesch 1976; Janis 1990; Marroig and Cheverud 2004; Mendoza et al. 2002) and interpretability in the context of either function or putative modules (see online table A2). Both datasets were corrected for variation due to subspecies and sex by adding to each value within a subsample the difference between the grand mean and the subsample mean (Sokal and Rohlf 1995; Marroig and Cheverud 2004). No significant effect of sex and subspecies was found on either the orientation or eccentricity of the covariance matrix for most species. Species for which I found a substantial effect, even if non-significant, were reduced to the largest single subspecies sample. A MANOVA test revealed no significant interaction between sex and subspecies for mean shape.

### Quantifying within-population covariation

A **P** matrix was calculated for each species with more than 27 specimens, including 2 species of tragulids, 13 species of cervids, and 32 species of bovids (see fig. 2 for details). Three subspecies of whitetailed and mule deer were analyzed separately, bringing the total number of cervid samples to 17. The sample size threshold of 27 was chosen based on a power analysis of rSDE (Haber 2011), with special consideration to include *Hydropotes inermis* due to its basal position. Variables were scaled by the species mean. Scaling by the mean mitigates the isometric effect of size variation among individuals (within and across species), as well as scale differences among variables, and ensures that the evolvability measures are meaningful in the context of Lande’s (1979) equation (Hansen et al. 2011). A non-weighted bending procedure was applied to each matrix to ensure that it is positive-definite (Jorjani et al. 2003; Pavlicev et al. 2009). Each sample was jackknifed specimen-by-specimen to identify aberrant specimens.

The average evolvability (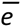), average conditional evolvability (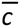), and average flexibility (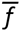) were calculated using simulations of random selection vectors (Marroig et al. 2009; Rolian 2009; Bolstad et al. 2014; see table 1). rSDE(**P**) was calculated based on the eigenvalues (see table 1) and adjusted for the effect of sample size by subtracting the mean rSDE(**P**) of a 1000 permutations from the observed value. This adjustment is equivalent to the correction suggested by Wagner (1984), but applicable to covariances as well as correlations. Confidence intervals were estimated using a non-parametric bootstrap procedure with a bias correction (BCa; DiCiccio and Efron 1996; Carpenter and Bithell 2000) and 999 iterations. The BCa correction was necessary because the pseudovalues distribution is expected to be biased upward and to depend on its mean when bounded by zero and/or one.

The evolution of rSDE(**P**), (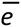), and (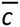) was explored by mapping these variables on the phylogeny and fitting several multi-optima Ornstein-Uhlenbeck (OU) models, as well as a single-rate Brownian motion (BM) model, in order to identify nodes where a shift in the clade’s typical parameter values has occurred (Butler and King 2004; Hipp and Escudero 2010). I tested for shifts in five nodes, using R package *maticce* (Hipp and Escudero 2010): all ruminants excluding tragulids (node number 8 in Hernández Fernández and Vrba 2005); Bovidae (node 15); Cervidae (node 25); Cervinae (node 27); and Caprinae (node 139). These nodes were chosen based on prior studies and a qualitative exploration of the data (figure 2; online figure A5). The comparison between bovids and cervids is the main focus of this paper, because of their different diversification patterns. Node 8FV was chosen in order to test the possibility that the difference between bovids and cervids is indeed due to a shift in either clades rather than basal to both (fully or partially). Caprinae and Cervinae were chosen because they seem to deviate in their typical eccentricity values from the rest of bovids and cervids, respectively. Caprines were found also to deviate in their integration pattern from all other bovids (Haber 2015). There was no a priori reason to test shifts in other nodes, and the observed patterns indicated that adding other models would only increase the number of models with effectively zero support. All possible combinations of the five putative transitions were tested, resulting in 32 alternatives of the multi-optima OU model and one single-rate BM model (see table 2). Variables were In-transformed for this analysis, and branch length was scaled by tree height. Models were selected based on the relative weights of their AICc scores, and the maximum likelihood estimate for each terminal taxon was calculated as an average of the 32 models, weighted by their AICc weights (Anderson et al. 2000).

**Table 2.** The different models describing the evolution of of rSDE(**P**) and 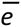. Based on the IL32 dataset. The best-supported models are in bold. K is the number of parameters in the model.

| Model No. | K | Nodes for which a shift is postulated <sup>b</sup> | | | | | $rSDE(P)$ | | $\bar{e}$ | |
| --- | --- | --- | --- | --- | --- | --- | --- | --- | --- | --- |
|  |  | 8FV<sup>a</sup> | Cervidae | Cervinae | Bovidae | Caprinae | AICc scores | AICc weights | AICc scores | AICc weights |
| 1 | 8 | 1 | 1 | 1 | 1 | 1 | 93.24 | 0.0096 | 11.11 | 0.0045 |
| 2 | 7 | 0 | 1 | 1 | 1 | 1 | 90.42 | 0.0392 | 8.28 | 0.0186 |
| 3 | 7 | 1 | 0 | 1 | 1 | 1 | 90.42 | 0.0392 | 8.28 | 0.0186 |
| 4 | 6 | 0 | 0 | 1 | 1 | 1 | 90.07 | 0.0466 | 6.08 | 0.0561 |
| 5 | 7 | 1 | 1 | 0 | 1 | 1 | 91.79 | 0.0197 | 8.31 | 0.0184 |
| 6 | 6 | 0 | 1 | 0 | 1 | 1 | 89.10 | 0.0759 | 5.62 | 0.0707 |
| 7 | 6 | 1 | 0 | 0 | 1 | 1 | 89.10 | 0.0759 | 5.62 | 0.0707 |
| 8 | 5 | 0 | 0 | 0 | 1 | 1 | 89.87 | 0.0517 | **3.50** | **0.2032** |
| 9 | 7 | 1 | 1 | 1 | 0 | 1 | 90.42 | 0.0392 | 8.28 | 0.0186 |
| 10 | 6 | 0 | 1 | 1 | 0 | 1 | **87.74** | **0.1497** | 8.51 | 0.0166 |
| 11 | 6 | 1 | 0 | 1 | 0 | 1 | 94.17 | 0.0060 | 10.12 | 0.0074 |
| 12 | 5 | 0 | 0 | 1 | 0 | 1 | 91.65 | 0.0212 | 9.35 | 0.0110 |
| 13 | 6 | 1 | 1 | 0 | 0 | 1 | 89.10 | 0.0759 | 5.62 | 0.0707 |
| 14 | 5 | 0 | 1 | 0 | 0 | 1 | **86.54** | **0.2728** | 5.96 | 0.0594 |
| 15 | 5 | 1 | 0 | 0 | 0 | 1 | 96.02 | 0.0024 | 8.69 | 0.0152 |
| 16 | 4 | 0 | 0 | 0 | 0 | 1 | 93.73 | 0.0075 | 7.81 | 0.0236 |
| 17 | 7 | 1 | 1 | 1 | 1 | 0 | 98.92 | 0.0006 | 12.79 | 0.0020 |
| 18 | 6 | 0 | 1 | 1 | 1 | 0 | 96.22 | 0.0022 | 10.09 | 0.0075 |
| 19 | 6 | 1 | 0 | 1 | 1 | 0 | 96.22 | 0.0022 | 10.09 | 0.0075 |
| 20 | 5 | 0 | 0 | 1 | 1 | 0 | 95.01 | 0.0039 | 7.97 | 0.0218 |
| 21 | 6 | 1 | 1 | 0 | 1 | 0 | 97.43 | 0.0012 | 10.12 | 0.0074 |
| 22 | 5 | 0 | 1 | 0 | 1 | 0 | 94.85 | 0.0043 | 7.54 | 0.0270 |
| 23 | 5 | 1 | 0 | 0 | 1 | 0 | 94.85 | 0.0043 | 7.54 | 0.0270 |
| 24 | 4 | 0 | 0 | 0 | 1 | 0 | 94.65 | 0.0047 | 5.51 | 0.0747 |
| 25 | 6 | 1 | 1 | 1 | 0 | 0 | 96.22 | 0.0022 | 10.09 | 0.0075 |
| 26 | 5 | 0 | 1 | 1 | 0 | 0 | 93.69 | 0.0076 | 9.58 | 0.0097 |
| 27 | 5 | 1 | 0 | 1 | 0 | 0 | 96.03 | 0.0024 | 10.18 | 0.0072 |
| 28 | 4 | 0 | 0 | 1 | 0 | 0 | 93.73 | 0.0075 | 9.15 | 0.0121 |
| 29 | 5 | 1 | 1 | 0 | 0 | 0 | 94.85 | 0.0043 | 7.54 | 0.0270 |
| 30 | 4 | 0 | 1 | 0 | 0 | 0 | 92.43 | 0.0144 | 7.14 | 0.0330 |
| 31 | 4 | 1 | 0 | 0 | 0 | 0 | 96.83 | 0.0016 | 8.50 | 0.0167 |
| 32 | 3 | 0 | 0 | 0 | 0 | 0 | 94.76 | 0.0045 | 7.43 | 0.0285 |
| BM | 2 | 0 | 0 | 0 | 0 | 0 | 102.01 | 0.0001 | 36.35 | 0.0000 |

<sup>a</sup> 8FV – Node No. 8 in Hernández Fernández and Vrba (2005), including Bovidae plus Cervidae without Tragulidae
<sup>b</sup> Nodes for which a shift is postulated are designated as 1. Nodes without a shift in the given model are 0

### Quantifying morphological diversification

Morphological diversification was measured by the overall amount, rate, and eccentricity of species dispersion in morphospace (table 1). These measures were calculated separately for each clade – Bovidae, Cervidae, Cervinae, and Caprinae, as above – based on its species means. Species means were scaled by the clade’s phylogenetically-weighted mean (the inferred root state, assuming a BM model within each clade, estimated using function *fastAnc* in the R package *phytools;* Revell 2012). The overall dispersion was calculated as the mean squared morphological distance between each species and the clade’s phylogenetically-weighted mean (equivalent to Foote 1993’s disparity). To facilitate comparisons, the expected distribution under a BM model was generated by simulating a 1000 datasets (function *sim.char* in the R package *Geiger;* Harmon et al. 2008) and recalculating disparity for each simulated dataset. These simulations also show that BM is a good enough approximation for evolution within each clade (see figure 3, top). The rate parameters for these simulations were provided by the clade’s evolutionary rate matrix (**D**_IC_ in table 1). **D**_IC_ is the covariance matrix of independent contrasts between species means, again assuming a BM model within each clade, and therefore represents the coevolution of traits while taking into account phylogeny (Revell 2007). The rate of dispersion is the trace of **D**_IC_. The eccentricity of dispersion is the relative standard deviation of the eigenvalues of **D**_IC_ (rSDE(**D**_IC_) in table 1). Thus, this metric measures how evenly species disperse in morphospace, and is the among-species equivalent of rSDE(**P**). Confidence intervals for rates and eccentricity of dispersion were estimated using a parametric bootstrap procedure with 500 replications assuming a multivariate normal distribution. Morphological diversification was also quantified directly from the landmark data, using Procrustes superimposition and PCA, yielding similar results as the interlandmark distances (online fig. A8).

### Evolvability in the direction of divergence

Species divergence was compared with the amount of mean-scaled phenotypic variance (evolvability) available in the direction of divergence, following Hansen and Houle (2008) and Hansen and Voje (2011). I first calculated the amount of variation that each species has along the eigenvectors of its clade’s evolutionary rate matrix (*e*(**d**_CL_); see table 1 for details), using its observed **P**. Species whose **P** matrices are more closely aligned with their clade’s diversification would have higher evolvabilities along the first eigenvector of **D**_IC_ than along other eigenvectors. I then calculated the amount of variation available for species in the direction of their own divergence from their clade’s inferred root state ((*e*(**d**_SP_) and *c*(**d**_SP_) in table 1; Hansen and Voje 2011), using the clade’s average **P** (**P**_AV_). Ideally, **P** here would be averaged over the clade’s history while accounting for phylogeny. However, previous analyses (Haber 2015) indicated that the phylogenetic structure of **P** within each clade is mostly random. Therefore, the simple average – i.e., assuming ‘white noise’ model – was preferred over other evolutionary models. The observed evolvabilities were compared to the expected evolvabilities in random directions (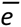 and 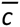; table 1), also calculated here from **P**_AV_ (Hansen and Houle 2008, Hansen and Voje 2011). Higher evolvabilities indicate species that have diverged closer to the direction of **P**_AV_ than expected by chance (Hansen and Voje 2011).

All analyses were repeated for three different phylogenetic hypotheses derived from the literature. Because results were essentially the same regardless of phylogeny, I present below results based on the phylogeny of Hernández Fernández and Vrba (2005) only. Results based on the other two phylogenies are presented in online appendix A. All analyses were carried out using R v.3.0.2 (R development Core Team 2013). The phylogenetic trees were manipulated using packages *ape* (Paradis et al. 2004) and *phytools* (Revell 2012). All R scripts, data, and phylogenetic hypotheses are available on Dryad (###).

## Results

### Within-population covariation

The two datasets (ILtes, IL32) yield largely the same results (online fig. A3), indicating that the smaller dataset captures most of the relevant information with its 32 variables. Matrix properties are consistently lower based on the IL32 dataset than those based on the ILtes dataset, and the signal is often weaker, probably due to a lower overall level of redundancy in the data. Adjusting rSDE(**P**) for sample size has little effect (online fig. A4).

Average flexibility (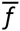) has a tight inverse correlation with the within-population eccentricity (rSDE(**P**)); average evolvability (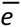) is positively but loosely correlated with rSDE(**P**); and average conditional evolvability (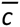) is not correlated with rSDE(**P**), nor with 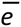 (fig. 1). 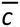 has yielded different results depending on shrinking tolerance, method of matrix inversion, and number of variables (online figure A3), and should therefore be regarded with caution. All rSDE(**P**) values deviate from zero after adjusting for sample size (fig. 2), indicating that all species deviate significantly from the null expectation of a random covariation structure. Bovids are in general less variable in their rSDE(**P**) than cervids (fig. 2); most bovid species do not differ from each other substantially, with the striking exception of Caprinae, whereas cervids show considerable variation throughout the clade and even among conspecific subspecies.

**Figure 1:**
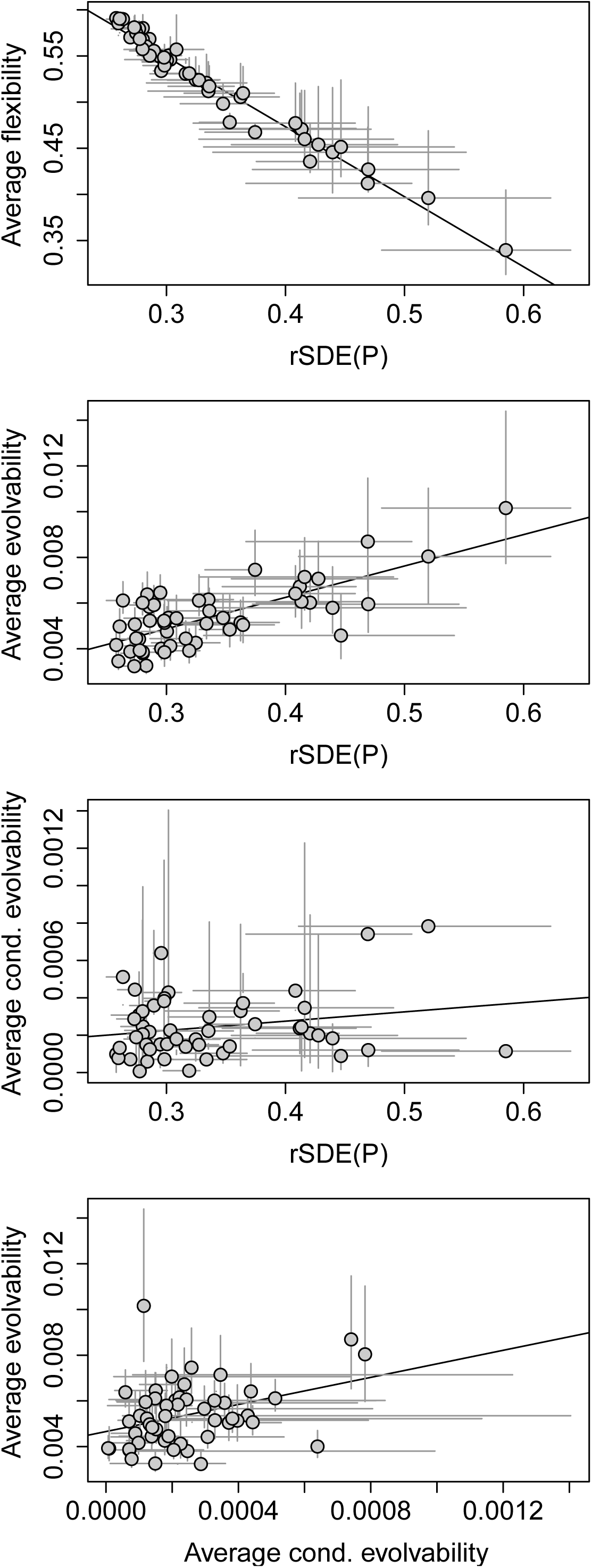
Comparison of **P**-matrix properties based on the IL32 dataset, mean-scaled. See table 1 for details on how these properties are measured and interpreted. Solid line indicates regression line. Grey bars indicate 95% confidence intervals using non-parametric bootstrap with BCa correction.

**Figure 2.**
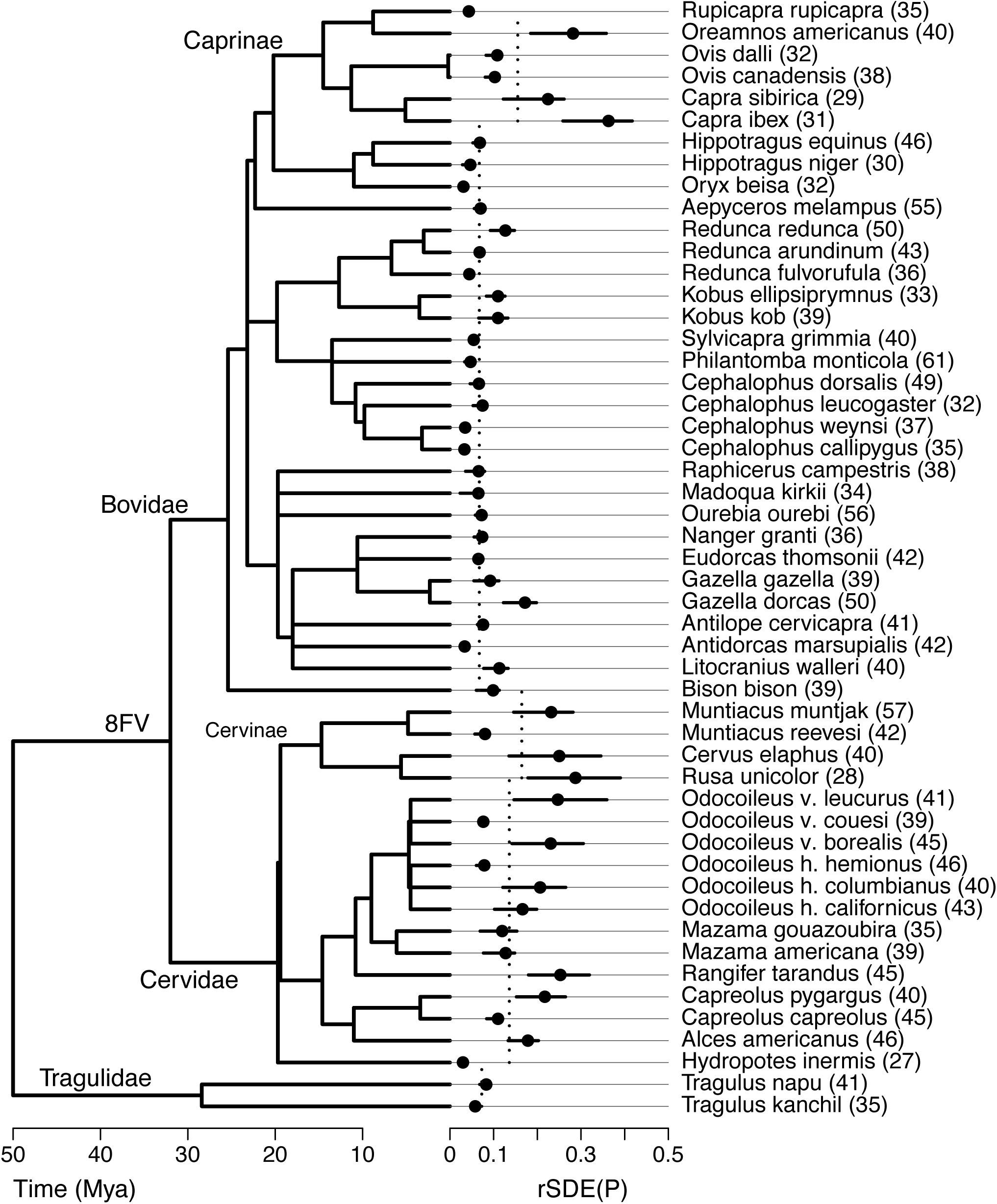
The relative standard deviation of eigenvalues (rSDE(**P**)) calculated from the IL32 dataset, mean-scaled. Values were adjusted for sample size by subtracting the mean permutation value from the observed value. Therefore, zero represents the expected value for a random matrix. Sample sizes are given in parentheses. Phylogenetic relationships follow Hernandez-Fernandez and Vrba (2005). Vertical dashed lines represent the maximum likelihood estimates (see table 3).

The model fitting results indicate that rSDE(**P**) and 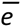 have not evolved following a Brownian motion process, but rather as a multi-optima OU process (table 2). The most strongly supported transition for rSDE(**P**) is between Caprinae and all other bovids; all models that do not include a shift at this node (models 17-32) have the lowest AICc weights. The best-supported model for all datasets (model 14 in table 2) includes a shift near the base of Cervidae as well. In addition, there is strong evidence against a transition that includes both bovids and cervids (node 8FV in table 2), as well as for all bovids. Therefore, bovids and cervids probably do not share the same typical rSDE(**P**) value. Bovids and tragulids, on the other hand, have the same typical value, which most likely characterized their ruminant ancestor as well. Model 10, which includes a shift near the base of cervines in addition to cervids and caprines, is also relatively well supported (see also online tables A3-A5). Therefore, there is some evidence that the typical rSDE(**P**) value of cervines is higher than that of other cervids. The best-supported model for 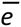 includes a transition for both bovids and caprines (model 8) and none for the other clades (table 2), implying that bovids were the ones to deviate from the ancestral ruminant state, rather than cervids.

Parameter estimates for the best-supported models are given in table 3, along with their weighted averages. The shifts inferred for rSDE(**P**) involve an increase for caprines and cervids (and possibly cervines). The shifts inferred for 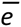 involve a decrease for bovids and an increase for caprines, relative to cervids and the ancestral state. The alpha estimates for rSDE(**P**) are high, yielding a phylogenetic half-life of only 8% of total tree height (calculated as log(2)/alpha; Hansen 1997), or 4 My out of the 50 My of ruminant history. The alpha and sigma-squared estimates are even higher for 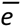.

**Table 3.**
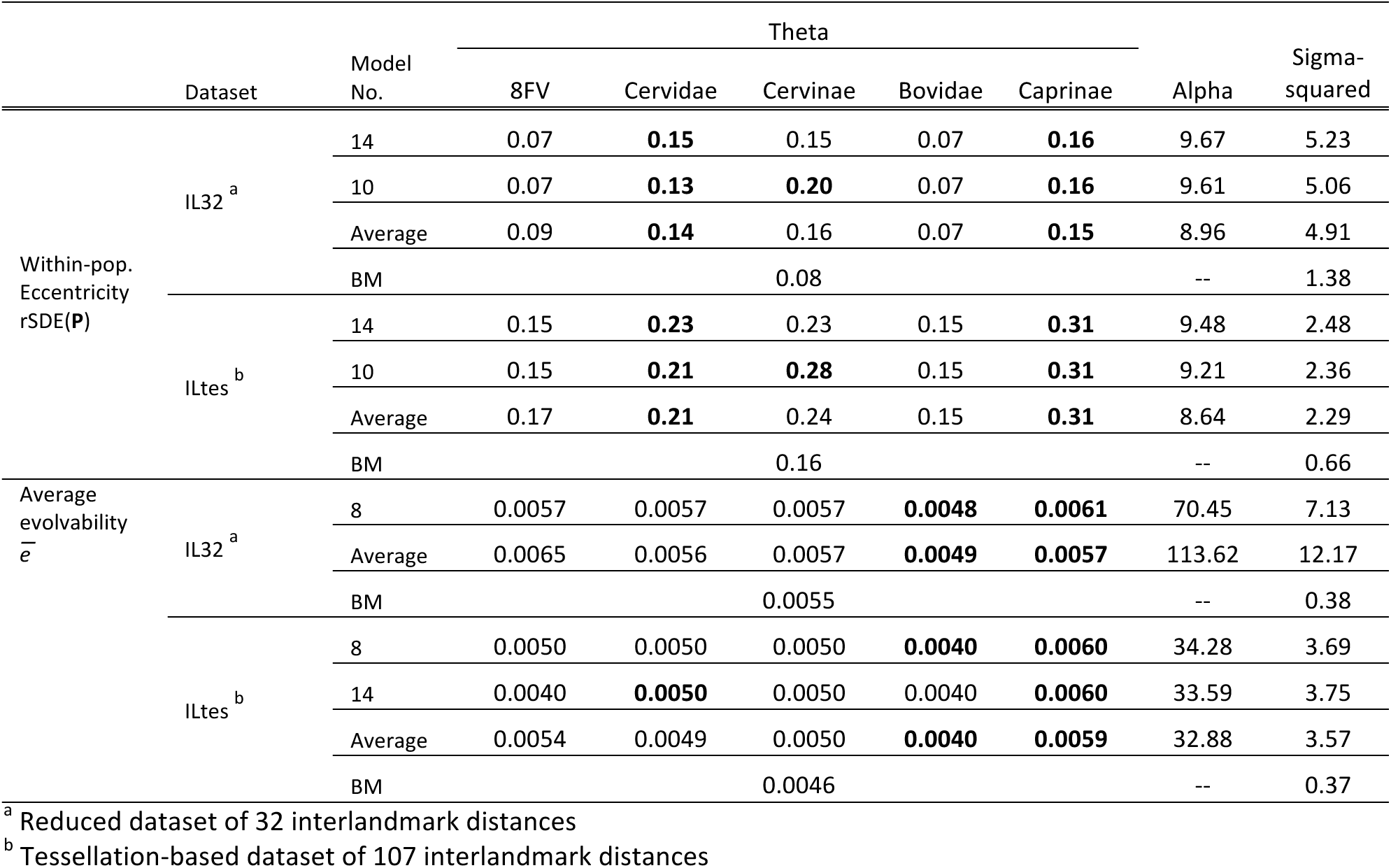
Maximum likelihood estimates for the best-supported models for the evolution of rSDE(**P**), and 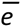 (see table 2 and online table A3). Nodes for which a transition was inferred are in bold. Estimates for a Brownian motion (BM) model are also given. Averages are weighted by AIC weights. All data are mean-scaled. Branch length is scaled by tree height. Theta values of rSDE(**P**) are adjusted for sample size as in fig. 1.

### Morphological diversification

The position of both species of *Alces* in morphospace deviates substantially from all other cervids (online fig. A2). Therefore, all measures were repeated for cervids with and without *Alces.* Clade disparity falls within the range expected under BM model for all clades (fig. 3, top), indicating that BM is a reasonable approximation for evolution within each clade. Bovids’ disparity is at the higher end of the expected range based on BM, especially when caprines are excluded, whereas cervids’ disparity is at the lower end, especially when *Alces* is excluded (fig. 3, top). Rates of dispersion are about the same for all clades (fig. 3, middle), but cervids’ eccentricity of dispersion, rSDE(**D**_IC_), is significantly higher than that of bovids, especially when *Alces* is excluded (fig. 3, bottom). Thus, bovids have dispersed further than cervids and into a wider range of directions. These findings are somewhat more pronounced based on ILtes and the Procrustes data, as well as the alternative phylogenies (online figures A6-A8). Caprines’ rSDE(**D**_IC_) and total disparity are lower than that of all bovids, and their rate of dispersion is higher (fig 3). However, based on the Procrustes data (online figure A8), caprine’s rSDE(**D**_IC_) is higher than that of all bovids.

**Figure 3.**
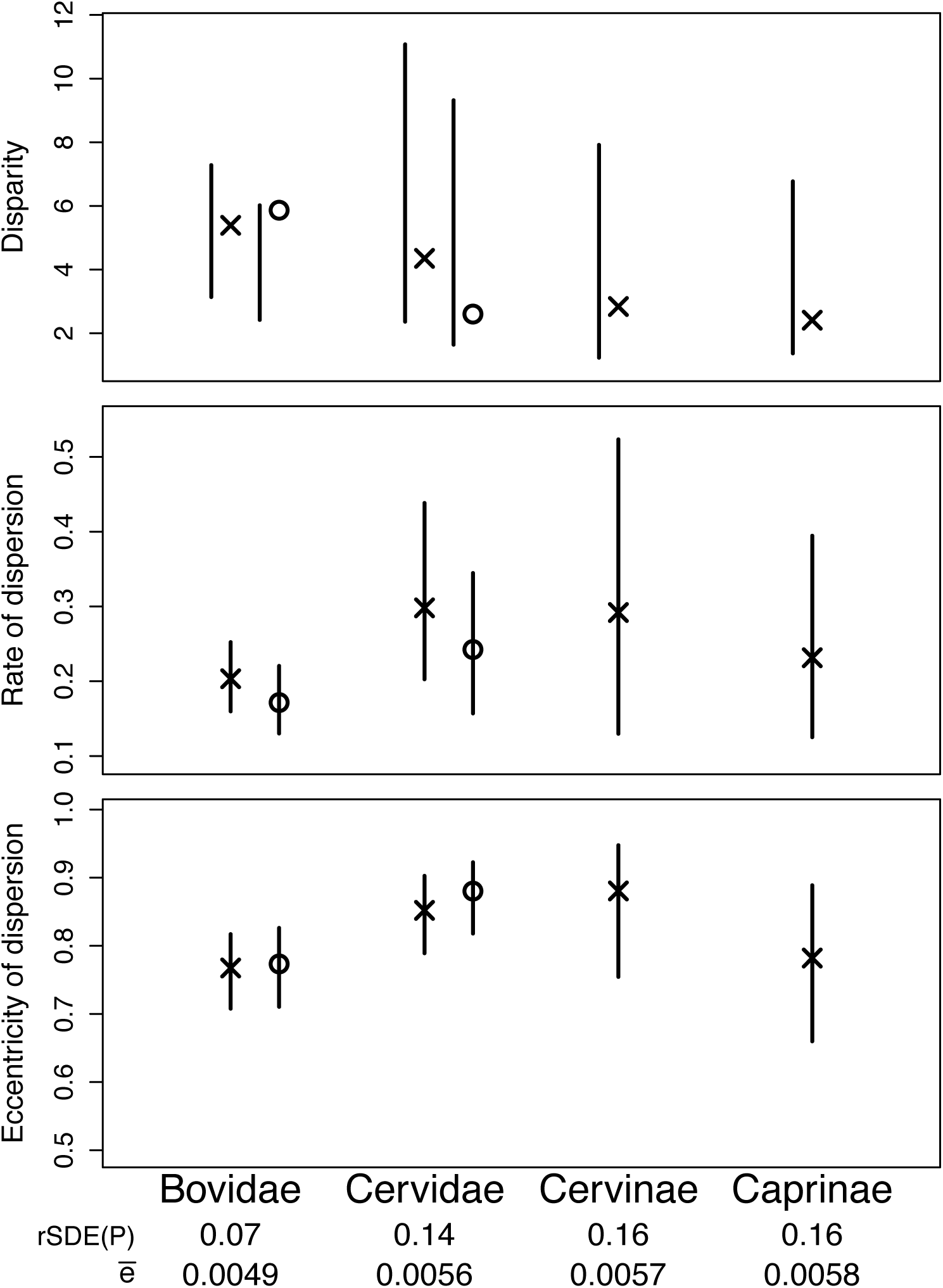
Measures of morphological diversification for the four clades that involve a shift in their typical within-population eccentricity (see and table 2). Maximum likelihood estimates of rSDE(**P**) and 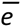 are also shown (see table 3). All metrics are proportional to the clade’s phylogenetically-weighted mean. Bars in the top panel (disparity) represent the expected distribution under Brownian motion model (95% interval). Bars in the lower two panels (rates and eccentricity) represent 95% confidence interval. Analyses involving Cervidae were repeated with (cross) and without (circle) *Alces.* Analyses involving Bovidae were repeated with (cross) and without (circle) Caprinae.

### Evolvability in the direction of divergence

Most species have substantially higher evolvabilities (*e*(**d**_CL_)) along the first eigenvector of their clade’s evolutionary rate matrix, **D**_IC_, than along other eigenvectors (fig. 4). The first eigenvector is associated with 76%-85% of the variation in **D**_IC_. The minimal overlap between the range of evovlabilities along the first eigenvector and those along other eigenvectors indicate that the vast majority of species have their **P** matrix more closely aligned with their **D**_IC_ than expected by chance. At the same time, the wide range of evolvabilities along the first eigenvector indicates that species differ greatly in their alignment. Cervids’ median value is only slightly higher than that of bovids, and their range is somewhat wider based on IL32 (though not based on ILtes; online fig. A9). Caprines have the same median and range as bovids. Excluding caprines from bovids (online fig. A11) and *Alces* from cervids (online fig. A12) made no difference.

**Figure 4:**
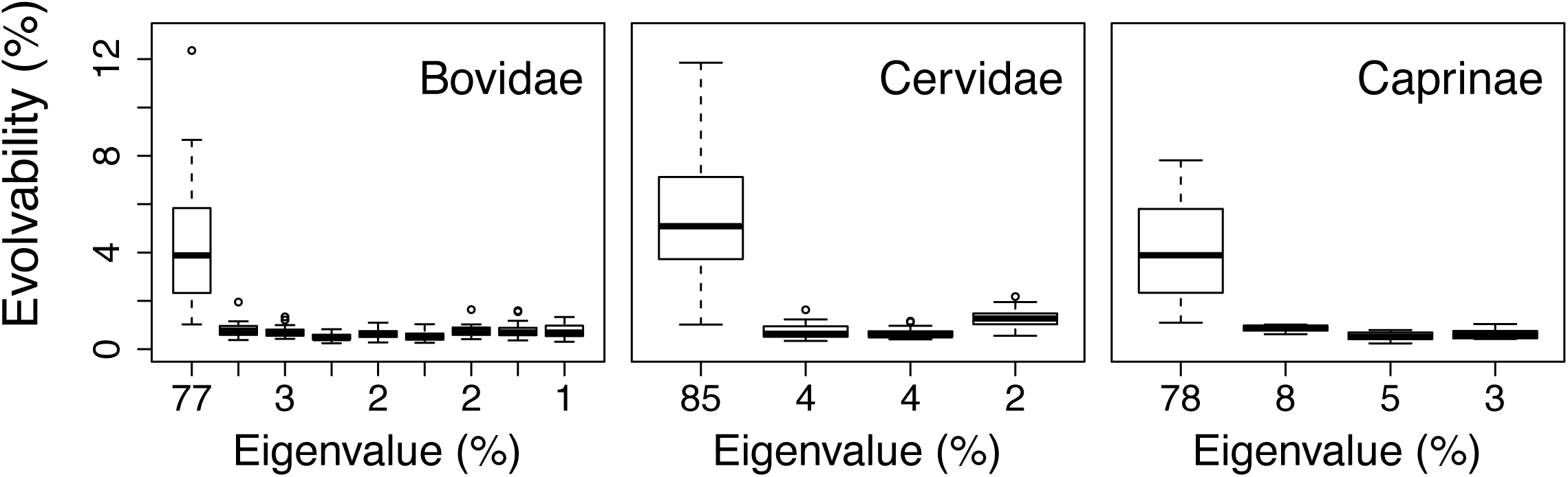
Evolvabilities in directions of clade’s dispersion (*e*(**d**_CL_)), measured as the amount of variation that each species has in its **P**-matrix in the direction of its clade’s eigenvectors. Eigenvectors are ordered by the relative size of their eigenvalues, indicated along the x-axis.

Most of the observed evolvabilities in the direction of species divergence – *e*(**d**_SP_) and *c*(**d**_sp_) – are closer to the maximum value than to the expected value for random directions (fig. 5). The average evolvability of **P**_AV_ is 0.31% of trait mean for bovids, 0.19% for cervids, and 0.06% for caprines, whereas the observed *e*(**d**_SP_) are above 0.4%, 0.6%, and 0.29%, respectively. The average conditional evolvability of **P**_AV_ is 0.08% for bovids, 0.05% for cervids, and 0.01% for caprines, whereas all the observed *c*(**d**_sp_) are an order of magnitude higher. Many of the bovid *c*(**d**_sp_) values are over 1%, whereas most cervid *c*(**d**_sp_) values are below 0.5%. When caprines are excluded from bovids, all bovid *c*(**d**_sp_) values are lower than 1% (online fig A17). When *Alces* is excluded from cervids, more *c*(**d**_sp_) values exceed 0.5% (online fig A18), thus narrowing the gap between bovids and cervids.

**Figure 5:**
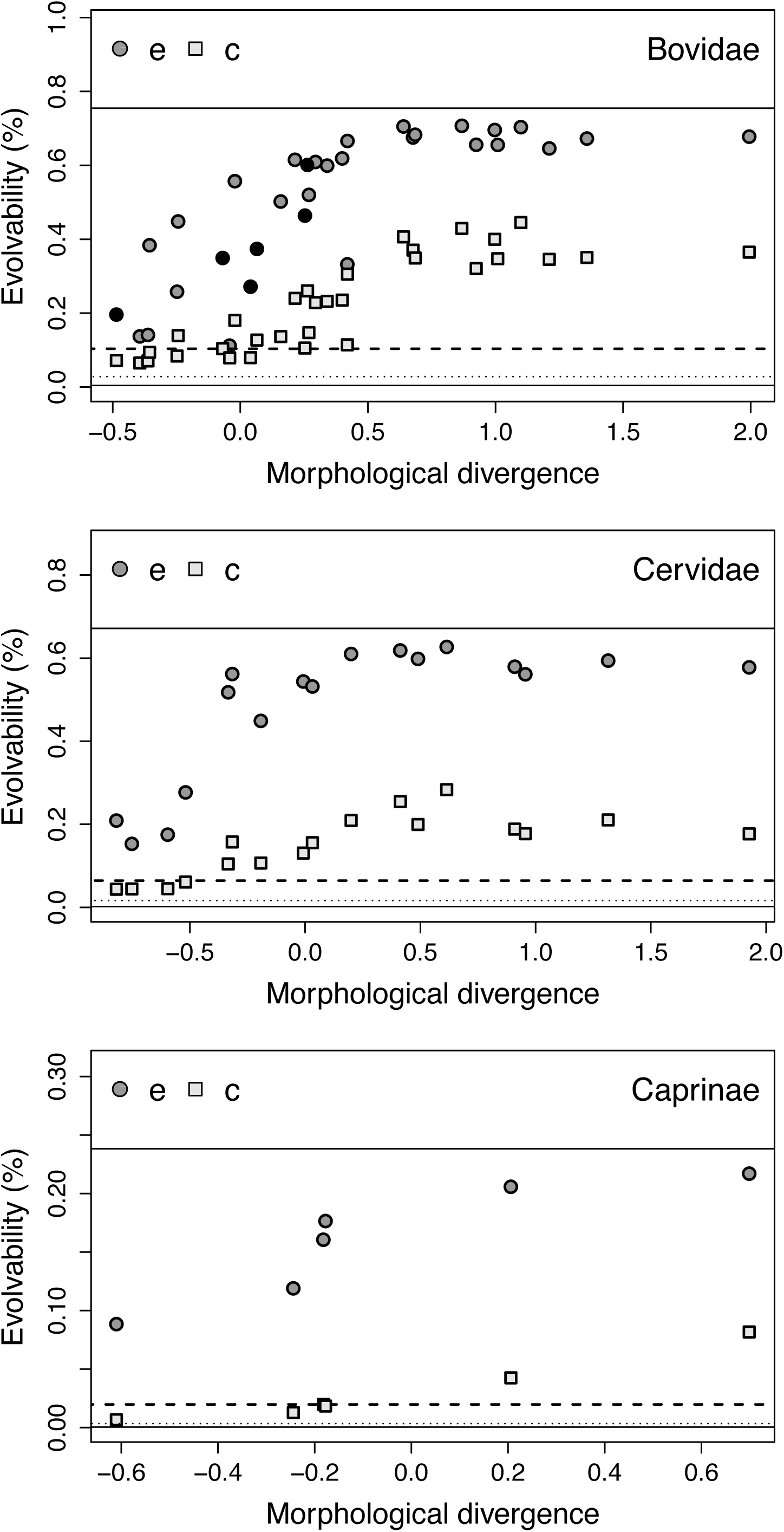
Evolvabilities (*e*(**d**_sp_)) and conditional evolvabilities (*c*(**d**_sp_)) in the direction of species divergence, calculated from the average within-species covariance matrix (**P**_AV_), plotted against the amount of divergence between each species and its clade’s ancestral state (*d*^2^; see table 1). Everything is proportional to trait mean. *d*^2^ is In-transformed for clarity. Horizontal bars mark the minimum and maximum evolvabilities (solid), the expected evolvability (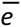, dashed) and conditional evolvability (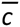, dotted). Caprines are highlighted in black in the top graph.

There is a distinct non-linear association between *e*(**d**_sp_) and *c*(**d**_sp_) and the magnitude of morphological divergence, *d*^2^ (fig. 5). Species whose divergence is more closely aligned with the direction of maximum evolvability managed to diverge further away from the clade’s ancestral state. The curve flattens close to the maximum value of evolvability at about 150% distance, in both bovids and cervids. Only species whose divergence is well aligned with the direction of the major axis of their **P**_AV_ (i.e., their observed *e*(**d**_SP_) value is close to the maximum) show divergence beyond that threshold.

## Discussion

The role of intrinsic constraints in determining macroevolutionary patterns is still an open question in evolutionary biology. The structure of variation and covariation within the population has been used for studying the role of intrinsic constraints on generational scale, but it is not yet clear which properties of that structure best predict evolutionary outcomes on the macro scale, and under what conditions. The question of why some clades are more diverse than others is of particular interest for macroevolutionary studies, and the contrast between bovids and cervids is a good example for that. Previous studies have provided ample evidence that bovids are substantially more diverse than cervids, taxonomically and ecologically (Allard et al. 1992; Spencer 1997; Sinclair 2000, Grubb 2002; Marcot 2004; Janis 2008; Heywood 2010). Based on discrete characters of teeth, Marcot (2004) has shown that bovids have been more diverse morphologically at least since the late Miocene (about 10 Mya). Here I provide additional evidence – based on their skull morphology – that bovids have a higher disparity than cervids and have dispersed into a wider range of directions in morphospace (fig. 3). So far, studies of the fossil record have not been able to explain these differences based on extrinsic factors such as environmental changes and biotic interactions (Janis 2008; Heywood 2010). My findings suggest that intrinsic constraints have played an influential role in the morphological diversification of ruminants, and particularly in the differences between bovids and cervids.

The role of intrinsic constraints was assessed here by comparing various properties of the within-population covariation (**P** and **P**_AV_; see table 1) with those of the among-population divergence (**D**_IC_), where **P** is assumed to reflect the potential to evolve and diversify and **D**_IC_ reflects the actual divergence that has occurred. Comparing **P** and **D**_IC_ in terms of their relative alignment reveals that most species have their **P** matrices more closely aligned with the major axis of their clade’s **D**_IC_ than with any other direction (fig. 4). Moreover, species whose divergence is more closely aligned with the direction of maximum evolvability have diverged further away from their clade’s ancestral state (fig. 5). In addition, cervids have lower evolvabilities along directions in which they have diversified less than bovids, and higher evolvabilities in directions in which they have diversified more than bovids (online fig. A13). Together, these results imply that intrinsic constraints have influenced the directions in which bovids and cervids have diversified. At the same time, there is a great variation in how well **P** matrices of different species are aligned with divergence, implying also a great variation among species in the orientation of their **P** matrices (fig. 4). This is in accord with a previous analysis (Haber 2015), which found that closely-related taxa differ in the orientation of their P matrices more than expected from their phylogenetic distance (assuming a single-rate BM model), and that bovid species differ from each other in their matrix orientation to the same degree as cervid species.

The positive signal in figures 4 and 5 is consistent with a role of intrinsic constraints in the morphological diversification of ruminants. However, without knowing the exact direction of selection throughout ruminant evolution, it is impossible to say whether the close alignment between **P** and **D**_IC_ is because covariation has biased divergence or because selection has been consistently aligned with covariation. That said, in the time span included here – 25 My for bovids and 20 My for cervids – it is not likely that selection would be pushing in the direction of **P** for that long unless **P** itself has aligned with selection, and such alignment would likely be due to its constraining effect (Jones et al. 2003; Arnold et al. 2008). Therefore, although not conclusive, these results suggest that the orientation and size of **P** have likely played a substantial role in determining the direction and magnitude of ruminant divergence. Thus, this study joins others (e.g., Hansen and Voje 2011) in demonstrating that even when covariance structure evolves relatively rapidly, intrinsic constraints could still bias the evolution and divergence of populations. At the same time, these analyses reveal little to no difference between bovids and cervids, and therefore do not provide a good explanation for why bovids are more diverse than cervids.

The analyses presented in figures 4 and 5 focus on the relative alignment between covariation and selection, accounting mostly for matrix orientation and size. As the time span and phylogenetic scale of the study increase, it becomes more difficult to reconstruct this relative alignment at any given time, and more likely that either or both have changed often enough that their relative alignment is no longer as informative or as relevant. Additional considerations might become more relevant for the macro scale, such as the heterogeneity of selection and the probabilities of extinction and speciation (Vermeij 1973; Liem 1973; Jablonski 2007; Gomulkiewicz and Houle 2009; Jones et al. 2012). Therefore, we might expect macroevolutionary patterns to be better predicted by properties that allow the population to respond effectively (i.e., reach close enough to the optimum to avoid extinction or competitive displacement) to a wide range of challenges. In other words, from a macroevolutionary perspective, it might be useful to look at the potential to respond to whatever may come, rather than the potential to respond to what has already come. Here, I considered four such measures: 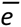, 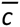, 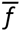, ‘and rSDE(**P**). The average conditional evolvability, 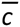, has yielded different results depending on shrinking tolerance, method of matrix inversion, and number of variables (online figure A3), due to its high sensitivity to sample size and matrix singularity. It is therefore considered less reliable in this study (although this is not necessarily the case for conditional evolvabilities along specific directions). The average flexibility, 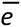, has a tight inverse correlation with rSDE(**P**) (fig. 1), in accord with expectations based on the number of variables and heterogeneity of the matrices (see online Appendix B). Therefore, as expected from the way they are computed, these two metrics capture effectively the same information regarding the potential to evolve into a wide range of directions. This further supports the interpretation of rSDE(**P**) as a measure of long-term flexibility. 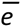 is positively but loosely correlated with rSDE(**P**) (fig. 1). Since rSDE(**P**) is scaled by matrix size, this result reflects more than just a scaling association between the mean and variance of the eigenvalues. Instead, it suggests that as matrix size increases, variance tends to be added more in directions with larger eigenvalues (thus increasing the variance of the eigenvalues) rather than randomly or evenly.

The model-fitting results indicate that these properties have evolved within a relatively limited range, which has shifted during the phylogenetic history of ruminants (fig. 2 and table 2). The main shifts supported by the data are near the base of cervids and caprines for rSDE(**P**), and bovids and caprines for 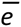 (online fig. A5). The high sigma-squared and alpha values (table 3) imply a great lability and little phylogenetic signal among closely-related taxa. Yet, the fact that the observed patterns are best explained by an Ornstein-Uhlenbeck model with multiple optima implies that these properties have been relatively constrained at the family and subfamily scale in spite of a great variation at the lower scales (Hansen 1997; Butler and King 2004). It could be argued that a good fit to OU model can also occur when a character evolves following Brownian motion with reflective boundaries (Harmon et al. 2010). rSDE(**P**) is indeed bound between 0 and 1, and 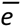 is bound by 0. However, this would not explain why they are bound within a different range for different clades. Therefore, these findings more likely suggest that rSDE(**P**) and 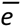 have been under selection, directly or indirectly, likely due to their association with morphological diversification.

The maximum likelihood estimates of rSDE(**P**) and 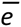 imply that bovids and cervids do not share the same typical value for either of them. However, the value of 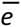 inferred for bovids is very close to that of cervids, and even slightly lower. Based on the current theory, we would expect the greater diversification of bovids to be associated with a higher average evolvability rather than a lower one, if constraints matter. Therefore, 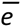 cannot explain the greater diversificaton of bovids. The rSDE(**P**) estimates, on the other hand, are easier to interpret here, especially in association with rSDE(**D**_IC_). The value of rSDE(**P**) inferred for bovids – as well as the ruminant ancestor – is lower than that of cervids (and independently, caprines; fig. 2 and table 3), suggesting a greater potential to respond effectively to a wider range of selective pressures. Accordingly, bovids have diversified into a wider range of directions in morphospace (a lower rSDE(**D**_IC_)), in addition to having a higher overall disparity (fig. 3). The finding that cervids have the same overall rate of evolution as bovids, even though their disparity is substantially lower (fig. 3), further suggests that cervids’ morphological evolution has been less diffusive than that of bovids, reverting back to previously-explored areas more often. Therefore, these findings support a scenario in which bovids have been able to take advantage of various ecological opportunities because they have retained the low within-population eccentricity (and higher flexibility) that they had inherited from their ancestor. Cervids, on the other hand, have taken the path of increasing eccentricity, which presumably has impeded their ability to take advantage of the same opportunities and has channeled their morphological diversification.

The results presented here support the idea that the long-term evolutionary success of lineages and clades – in terms of their morphological diversification at least – is influenced by their ability to respond effectively to a wide range of selective pressures (Vermeij 1973; Liem 1973; Draghi and Wagner 2008). In addition, this study suggests that the best predictor of that ability, at least in this case, is their within-population matrix shape, measured as rSDE(**P**) or average flexibility. More empirical and theoretical work is required in order to establish the universality of these findings and the validity of the assumptions underlying this association. Some relevant insights come from simulations by Draghi and Wagner (2008), who show that populations whose distribution of mutational effects is less eccentric can adapt faster to a wider range of circumstances. Simulations by Goswami et al. (2014) also suggest that clades with lower rSDE(**P**) disperse more evenly in morphospace and have somewhat higher disparity (although evolution was modeled here as BM so heterogeneity of selection is not incorporated). Some empirical evidence comes from research on morphological integration, assuming rSDE(**P**) reflects the magnitude of integration. For example, lower integration was found to be associated with greater functional divergence in great apes (Rolian 2009; Rolian et al. 2010; Grabowski et al. 2011) and higher module disparity in Carnivora (Goswami and Polly 2010). In addition, Claverie et al. (2013) found that modules that evolve more independently tend to evolve faster in mantis shrimp, although they did not look at the within-population level directly.

Several assumptions underlie the suggested association between matrix shape and macroevolution. The main assumption is that selection changes frequently enough and in directions different enough from the major axis of **P** (or **G**), thus providing the opportunities for testing and realizing the potential reflected in matrix shape. This is ultimately an empirical question, and some evidence already implies that selective pressures can be highly dynamic at various scales and dimensionality (Estes and Arnold 2007; Uyeda et al. 2011; Siepielski et al. 2009). A study of fossil sequences that span multiple episodes of climate change found a consistent decrease in rSDE(**P**) for two carnivoran lineages associated with these episodes (Goswami et al. 2015). However, with increasing concerns over global climate change, understanding and predicting population response to erratic environments becomes increasingly relevant regardless of whether this has been common in the past or not. Simulations could reveal how heterogeneous selection needs to be in order for the potential to respond to a wide range of selective pressures to become more relevant than the potential to respond faster in any particular direction.

A second assumption is that there is a threshold of variance that needs to be exceeded in order for the population to persist long enough and respond effectively. Theoretical work and simulations show that stochastic movements of the optimum increase the chances that a population will drop below its carrying capacity (Arnold et al. 2001, Gomulkiewicz and Houle 2009, Jones et al. 2012, Kopp and Matuszewski 2014), and that higher eccentricity exacerbates this effect because it increases the chance of not having enough variation in the desired direction (Jones et al. 2012). Yet another assumption is that matrix shape remains effectively stable long enough so that **P** and **G** can affect evolution through their eccentricity even when their orientation and size change. This, again, is an empirical question for which there is currently too little data. For the ruminant skull, there is some evidence that matrix shape is more stable than matrix orientation: pairwise comparisons of matrix orientation across ruminant species vary within 50% of their possible range (0.4-0.9; Haber 2015), whereas eccentricity varies within 33% of its range (figs. 1 and 2). Similar numbers were found for covariance matrices of the fruit fly wing when changes are induced by mutations (Haber and Dworkin 2015). Simulations by Jones et al. (2003, 2012) also indicate that eccentricity tends to vary less than orientation for most combinations of genetic architecture and selection.

The association found here between matrix shape and macroevolutionary patterns suggests a trade-off between the ability to evolve into a wide range of directions – i.e., long-term flexibility – and the ability to evolve faster along a particular direction – i.e., short-term robustness – as was also suggested by Goswami et al.’s (2014) simulations. This, in turn, suggests a possible dissociation between macro- and microevolution, where clade dispersion on the macro scale is largely associated with matrix shape and the heterogeneity of selection, while population divergence on the generational scale is largely associated with the relative orientation and magnitude of covariation and selection. Again, comprehensive simulations could reveal the range of parameters and conditions in which different properties become more relevant. In addition, matrix shape could be linked to macroevolution through taxonomic diversification, if the probability of reproductive isolation increases when the population moves further away from the major axis of covariation, regardless of how far it moves. This would be in accord with other studies that point to a decoupling between population-level processes and macroevolutionary patterns (Jablonski 2007; Rabosky and Matute 2013). Although the focus here is on macroevolutionary patterns, the same rationale should hold for short-term evolution in highly dynamic environments, an issue that has become increasingly relevant with the advance of global climate change. Therefore, studying variational properties that reflect the potential to respond to a wide range of selective pressures – and promote long-term flexibility – could be useful for understanding population response to erratic environments as well as macroevolutionary patterns.

## Acknowledgements

I owe many thanks to my advisors and committee members for their invaluable support, guidance, and advice: Mark Webster, Leigh Van Valen, David Jablonski, Michael Foote, Peter Wagner, Ian Dworkin, Jeff Connor, and Tamar Dayan. This study has benefited greatly from discussions with Miriam Zelditch, Thomas Hansen, Gabriel Marroig, Mihaela Pavlicev, David Bapst, and two anonymous reviewers, and their thoughtful comments. I thank the following people and institutions for access to their collections and databases: D. Wilson and L. Gordon, The Smithsonian Institution; M. Schulenberg, Field Museum of Natural History; E. Westwig, American Museum of Natural History; R. Sabin, Natural History Museum, London UK; M. Herman, Quex Museum; T. Sharib, Tel Aviv University; F. Mayer, Museum für Naturkunde, Berlin DE; H. van Grouw, Naturalis; C. Conroy, Museum of Vertebrate Zoology at Berkeley; M. Flannery, California Academy of Sciences; J. Chupaska, Museum of Comparative Zoology at Harvard; J. Dines, Natural History Museum of Los Angeles County. Funding for this project was provided by the National Science Foundation (DDIG DEB-0709750), University of Chicago Hinds Fund, University of Chicago Women’s Board Travel Award, AMNH Collection Study Grant, Dan David Fellowship, Center for Absorption in Science at the Ministry of Absorption, Israel, Council of Higher Education, Israel, and BEACON Center for the Study of Evolution in Action at Michigan State University. This material is based in part upon work supported by the National Science Foundation under Cooperative Agreement No. DBI-0939454. Any opinions, findings, and conclusions or recommendations expressed in this material are those of the author and do not necessarily reflect the views of the National Science Foundation.

